# Cell-level random splits leak group-owned answers in single-cell benchmarks

**DOI:** 10.64898/2026.09.09.750484

**Authors:** Hu Cang, Sha Sun

**Affiliations:** Department of Developmental and Cell Biology, University of California, Irvine, Irvine, CA 92697, USA

## Abstract

Machine learning models in single-cell biology increasingly forecast differentiation, reprogramming and therapeutic response from early transcriptomic profiles. Testing whether a model has learned real biology requires held-out cells. Single-cell data, however, are grouped: cells from the same clone, patient or batch share the same label. A random split therefore places relatives of each test cell, carrying its label, in the training set, and a model can score well by memorizing a relative instead of learning a transferable rule. Grouped validation removes this leakage but leaves far fewer independent units behind each error bar. Here we show how to estimate this leakage before training any model, from two properties of the data: exposure, the fraction of test cells with relatives in training, and retrievability, how often a nearest-neighbor search returns such a relative rather than an unrelated cell. Across lineage-barcoded and patient data, exposure determines whether a random split opens a leakage channel, and retrievability determines how much it can inflate the score. The inflation is negligible where cell state has decoupled from ancestry, much larger where clonal sisters remain close in expression space, and in a patient cohort large enough to overturn a clinical conclusion. We also provide `leakcheck`, which computes both properties in seconds, before the outcome model is fitted.

## Introduction

As single-cell genomics transitions from descriptive atlases to predictive machine learning, computational models increasingly forecast cellular differentiation, reprogramming, and therapeutic response directly from early transcriptomes [1, 2, 3, 4, 5, 6, 7, 8, 9, 10]. Evaluating whether these models learn transferable biology requires held-out validation. Standard cross-validation assumes independent and identically distributed (*i*.*i*.*d*.) test observations [11, 12, 13, 14, 15], yet single-cell measurements routinely violate this assumption. Single-cell datasets possess hierarchical group structures: cells share progenitor clones, clinical biopsies originate from individual donors, organoids share culture wells, and libraries introduce batch effects [16, 17, 18, 19, 20, 21, 22]. In each setting, an underlying biological unit owns the outcome label and shares it across many cells.

Random cross-validation scatters members of the same biological group across training and test folds. Target labels of held-out cells therefore reside in training, attached to clonal or patient relatives. A predictive model can score well simply by locating a training relative and copying its label rather than learning a transferable biological rule. Random splitting turns generalization testing into an open-book exam: answers to held-out queries already sit in the training data.

The established remedy is grouped validation: holding out entire clones, donors, or batches intact [12, 13]. Grouping closes this shortcut and aligns validation with independent biological units, but be-cause single-cell cohorts contain far fewer biological groups than cells, grouping widens fold-to-fold variance and alters effective degrees of freedom.

Methodological practice remains divided: multiple-instance learning architectures treat donors as intact units by construction [23, 24, 25, 26, 27, 28, 29, 30, 31], and leading foundation models partition across donors [1, 2, 32], yet standard machine learning splitters default to cell-level partitioning unless grouping keys are explicitly supplied [33], and recent audits note that benchmarks reflect whatever their setup permits [34, 35].

Here we show that two pre-training properties bound how much leakage a split can produce: exposure (*ε*), the proportion of test cells with relatives in training (0 ≤ *ε* ≤ 1), and retrievability (*r*), the frequency with which a nearest-neighbor query in expression space retrieves a true relative rather than an unrelated cell. Exposure determines whether a leakage channel exists; retrievability determines how easily an algorithm accesses that channel in transcriptomic space. Their product *εr* represents the proportion of test cells for which the channel is open and accessible.

To calibrate these properties, we analyzed three expressed DNA lineage-barcoding systems where heritable barcodes identify clonal relatives unambiguously (*y_i_* = *y*_clone(*i*)_): hematopoietic differentiation (LARRY [36]), direct transcription-factor reprogramming (CellTag-multi [37]), and drug tolerance in lung adenocarcinoma (PC9 [38, 39, 40]). We then extended the framework to a clinical cohort where labels belong to human donors: dementia in the Seattle Alzheimer’s Disease Brain Cell Atlas (SEA-AD) [41].

Across all three lineage systems, we evaluated a panel of six supervised architectures spanning standard practice [33, 42, 43, 44]: an uninformative prior classifier, uniform *k*-nearest neighbors, tuned *L*_2_ logistic regression (employed by CellRank [7]), radial basis function (RBF) Nyström kernel logistic regression, gradient-boosted decision trees (among architectures compared by CellTag-multi [37]), and a tuned multilayer perceptron [9, 23]. Scoring each model under both cell-level random splitting and clone-grouped holdout defines the evaluation gap: Δ*L* = *L*_grouped_ − *L*_random_.

We demonstrate, first, that exposure alone cannot predict empirical inflation: all three lineage systems are heavily exposed (*ε* = 0.483, 0.718, and 0.943 in LARRY, CellTag, and PC9), yet PC9 inflates least and CellTag most (0.102 versus 0.0032 nats), inverting the exposure ranking. Second, retrievability in gene expression space tracks realized leakage: surviving cancer persisters occupy transcriptomic states where ancestral lineage no longer predicts nearest neighbors (*r* = 0.0084 versus *r* = 0.435 in CellTag), and this geometric retrieval orders score inflation across all six architectures (CellTag > LARRY > PC9). Third, in a clinical cohort, donor-level scoring resolves whether this gap alters scientific conclusions: in dementia, holding out donors changes the clinical conclusion (74 of 85 donors called correctly under random splitting versus 59 under grouped validation), demonstrating that random cross-validation can produce an illusion of diagnostic readiness. Finally, we release `leakcheck`, an open-source tool that computes *ε*, *r*, and hierarchical group structures in seconds before model training.

## Results

### Clonal exposure under random splitting is predictable from clone sizes alone

To test how biological hierarchy affects validation independence, we analyzed three expressed lineage-barcoding systems (Fig. 1a,b). In each system, progenitor cells were tagged with heritable DNA barcodes, early transcriptomes were profiled by single-cell RNA sequencing (scRNA-seq), and mature descendants were cultured onward to determine cell fate. Because clonal relatives inherit the same barcode, sister cells carry bit-identical labels (*y_i_* = *y*_clone(*i*)_). When cells are partitioned into *K*-fold cross-validation at random, clonal relatives scatter across training and test folds. We define clonal exposure, *ε*, as the proportion of held-out test cells with at least one clonal relative in training, computable from clone sizes and fold count alone before inspecting gene expression.

**Figure 1:**
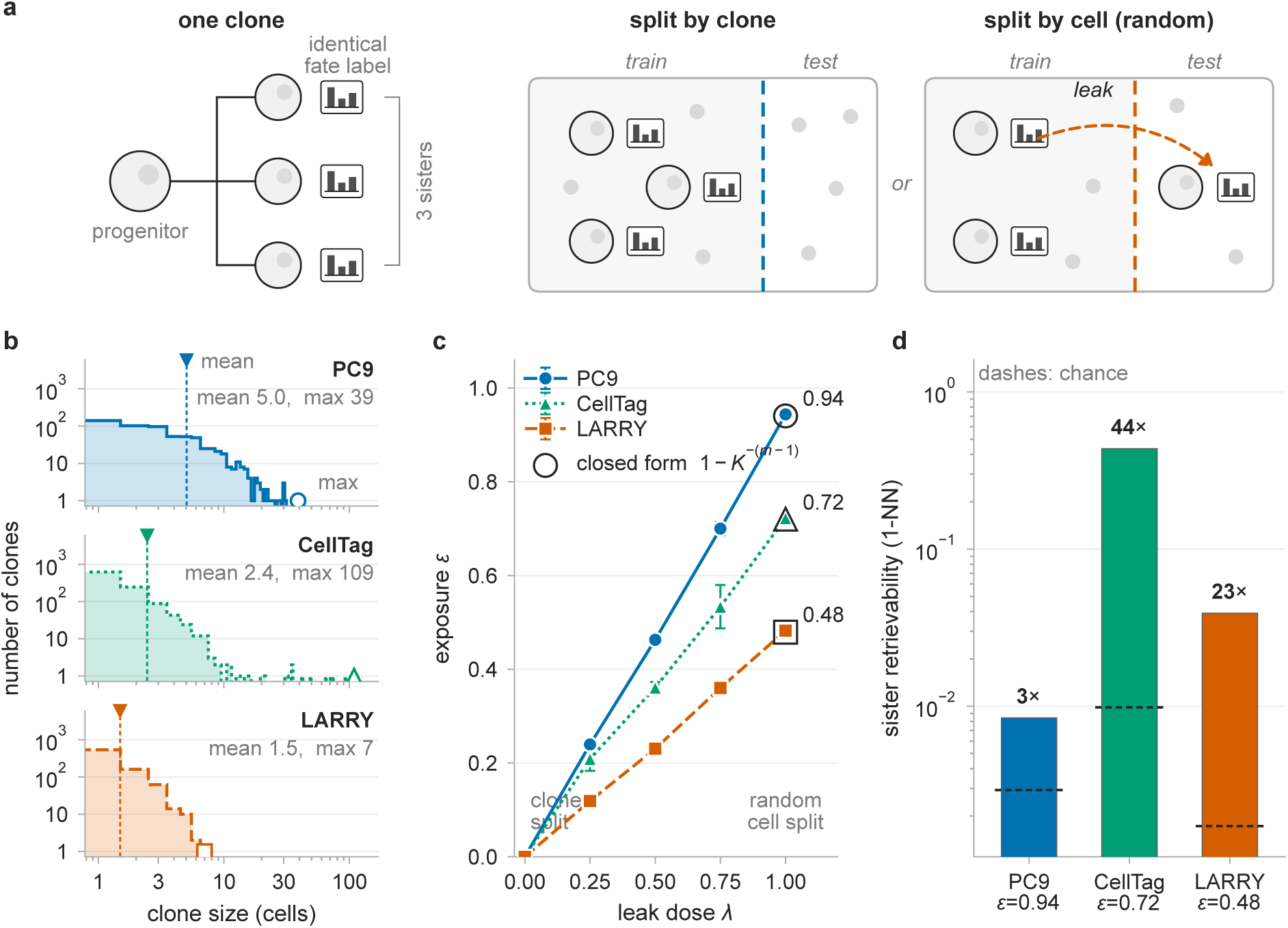
A clone owns the label, and a cell-level random split puts copies of it on both sides of the partition. **a**, In a lineage-barcoding experiment the fate outcome is measured on mature descendants and assigned to every early sister sharing the barcode, so sisters have a bit-identical fate vector *y*_*i*_ = *y*_clone(*i*)_. Clone-grouped cross-validation holds whole clones out and closes the channel. Cell-level random cross-validation scatters sisters across the partition, which allows a test cell to be matched to a training relative carrying its own label. **b**, Distribution of profiled cells per clone in each system, with mean and maximum annotated; note the log axes and the 109-cell tail in CellTag. **c**, Clonal exposure against leakage dose λ, the fraction of multi-cell clones dissolved into cell-level units before fold assignment, drawn as nested subsets so that the sweep is a titration rather than five unrelated draws; λ = 0 is a clone-grouped split and λ = 1 a cell-level random split. Filled markers are the mean over three seeds and error bars their standard deviation; open markers at 1 are the closed form *ε* = 1 − *K*^−(*m*−1)^ averaged over cells, each clone weighted by the number it contributes, and the two agree to within 0.0043. Exposure can be computed from clone IDs alone, before any model is fitted. **d**, Nearest-neighbor sister retrieval rate on a logarithmic scale. Bars are measured rates, dashed lines the analytical chance level for each system, labels the fold-enrichment over chance. PC9 has the highest exposure of the three (*ε* = 0 943, panel c) and the lowest retrievability (*r* = 0 0084): counting relatives ranks the systems PC9, CellTag, LARRY, whereas finding them ranks the systems CellTag, LARRY, PC9, the order in which they inflate.

The three cohorts span divergent clone-size architectures (Fig. 1a,b). LARRY hematopoiesis (1,167 day-2 cells across 785 clones) features compact, uniformly sized clones (mean 1.49 cells, maximum 7) differentiating into 6 blood lineages [36]. CellTag direct reprogramming (2,606 day-3 cells across 1,066 clones) carries a heavy clone-size tail (mean 2.44, maximum 109) transitioning toward induced endoderm progenitors [37]. PC9 lung adenocarcinoma (3,280 day-3 cells across 651 lineages) exhibits deep clonal expansion (mean 5.04, maximum 39) under osimertinib selection into drug-tolerant persisters [38].

Under 5-fold random cross-validation, clonal exposure reaches *ε* = 0.483 in LARRY, 0.718 in CellTag, and 0.943 in PC9 (Fig. 1c), whereas clone-grouped splitting sets *ε* = 0 by construction. For a clone of size *m* partitioned uniformly across *K* folds, the probability that a held-out cell has at least one training relative is

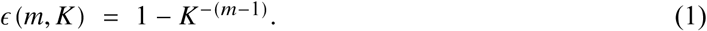

Weighting each clone by cell count yields closed-form predictions of 0.479 for LARRY, 0.720 for CellTag, and 0.940 for PC9, agreeing with empirical fold assignments within 0.0043 (Methods).

For singleton clones (*m* = 1), exposure is zero, whereas for multi-cell clones, *K*^−(*m*−1)^ decays exponentially, causing *ε* to saturate rapidly. Exposure reflects the proportion of cells in multi-cell clones rather than maximum clone size: CellTag carries the heaviest clone-size tail (109 cells) yet is less exposed than PC9 (39 cells). For experimental design, shallowly barcoding many clones minimizes exposure, whereas deep clonal expansion maximizes it. Exposure opens the leakage channel; we next evaluate its theoretical capacity before examining expression-space retrieval (*r*; Fig. 1d).

### An unparameterized oracle bounds channel capacity

To quantify the theoretical ceiling of this leakage channel, we constructed a zero-parameter sister-lookup oracle. For held-out cells with clonal sisters in training, the oracle copies that sister’s mature fate vector; for unexposed cells, it predicts the training base rate. The oracle reads no gene expression, fits no parameters, and learns no rules. Performance is measured by out-of-fold cross-entropy loss in nats (lower is better; a 0.1-nat reduction assigns ∼ 10 % more probability to the true fate). An uninformative prior classifier predicting base rates achieves 1.379 nats on LARRY as reference baseline, while tuned multinomial logistic regression on 50 principal components evaluated under clone-grouped holdout serves as the generalization comparator (0.914 nats, evaluated within the oracle experiment’s own fold draws and therefore slightly different from the Table 1 entry; Methods).

**Table 1:**
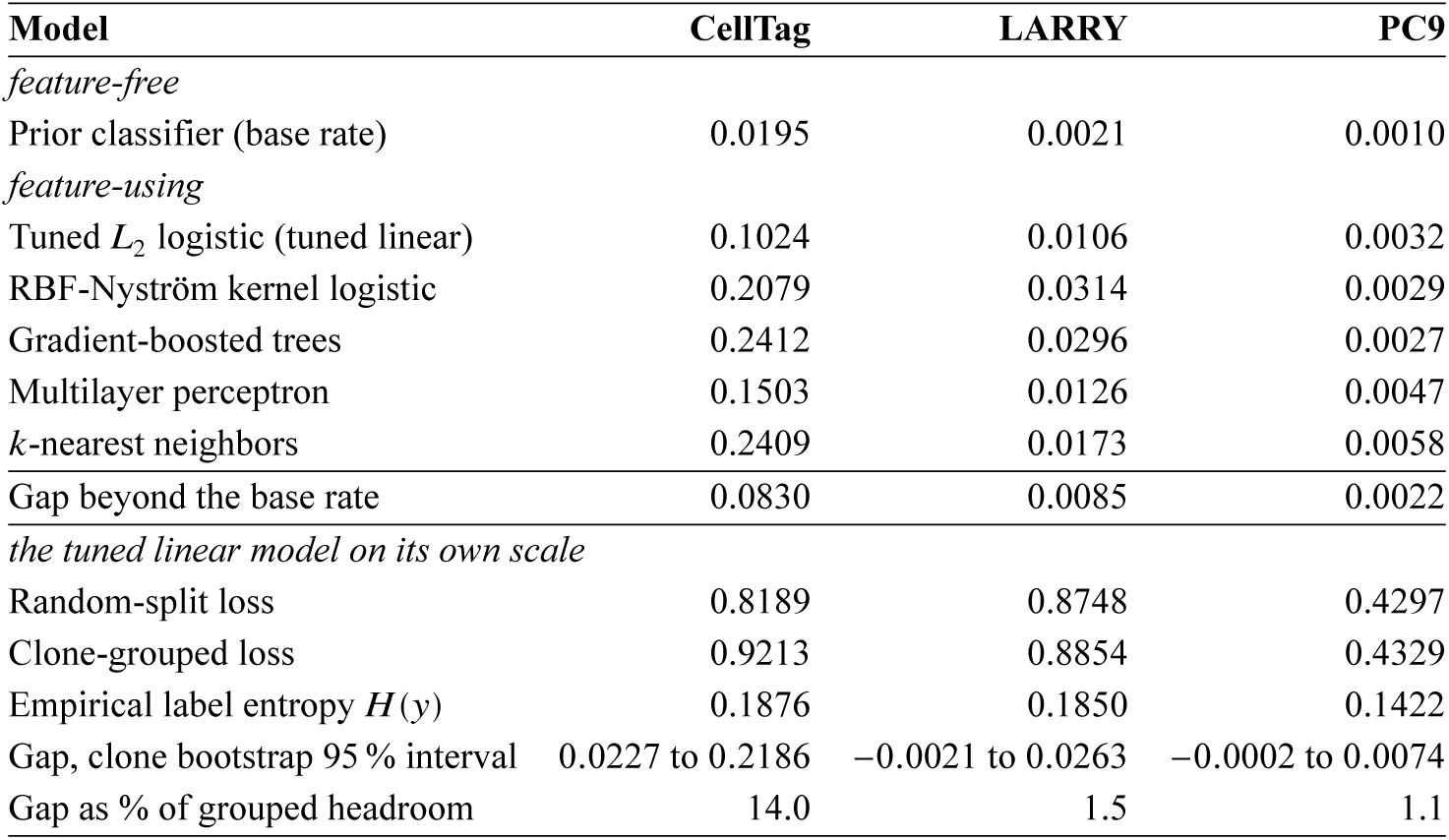
Evaluation gap Δ*L*_grouped_ − *L*_random_ in nats for each of the six models. All six models run on the same cells, features and fold geometry, with the split as the only variable. The first model, the prior classifier, predicts the training-fold base rate and reads no gene expression, so its movement is feature-free and cannot be mediated by neighbor retrieval; the row *Gap beyond the base rate* is the tuned linear gap net of it. The ordering CellTag LARRY PC9 holds on the point estimates for every model and survives that subtraction. Every gap is a mean over 3 split seeds of the cell-weighted out-of-fold loss; every model is group-aware throughout, including the multilayer perceptron’s early-stopping holdout (Methods give the cell-level value). The interval on the tuned linear model is a percentile bootstrap over 2,000 draws that resamples clones, not cells. Grouped headroom is the clone-grouped loss less the empirical label entropy, which places tasks of different class count and entropy on one scale.

This zero-parameter oracle outperforms the trained linear model (Fig. 2a). Under 5-fold random cross-validation, the oracle achieves an out-of-fold loss of 0.844 ± 0.017 nats across 5 seeds, surpassing the clone-grouped linear model by 0.070 nats and assigning 7.2 % more probability to the true fate. Under clone-grouped splitting where sisters are absent, the oracle returns to 1.379 nats (1.63 times higher), confirming its advantage stems entirely from leakage. On exposed cells alone (0.477 of cells), the oracle reaches 0.195 nats versus 0.949 nats for the grouped linear model (4.86 times lower), matching the entropy of the cell’s own empirical fate vector.

**Figure 2:**
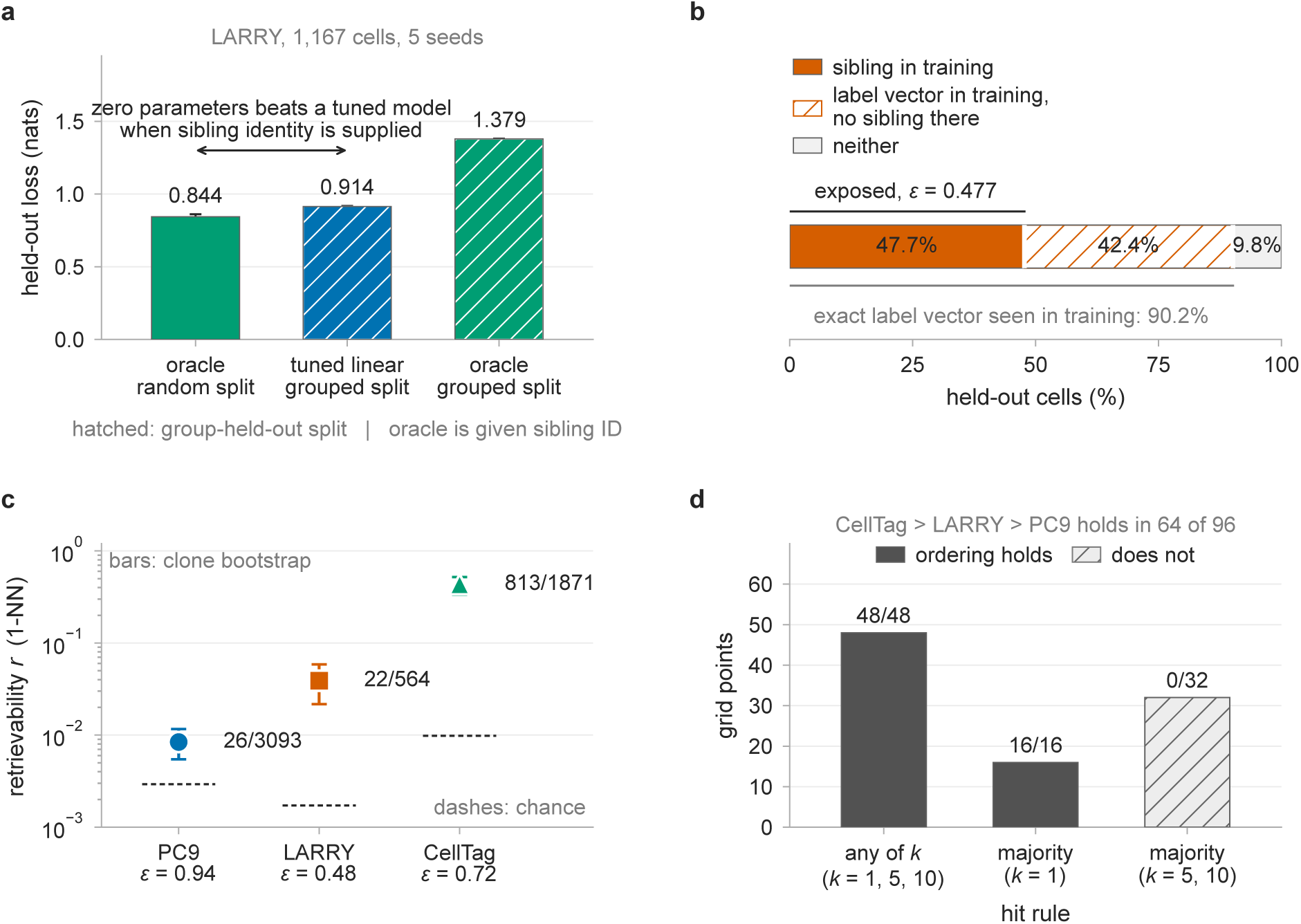
Channel capacity and expression-space retrieval fidelity. **a**, The sister-lookup oracle on LARRY, 1,167 cells, mean ± SD over 5 seeds. The comparison is deliberately asymmetric: the oracle is given sister identity and reads no expression at all, whereas the trained linear model must find its relatives in expression space. Under a random split a rule with no parameters and no features scores 0.844 nats, better than the trained linear model evaluated under a clone-grouped split (0.914 nats). The same oracle under a clone-grouped split has no sister to copy and falls back to the base rate (1.379 nats, the right-hand bar; hatching marks a group-held-out split). The bars quantify the leakage channel; because the oracle returns the base rate wherever a sister is absent, they do not bound a model that also learns transferable biology. **b**, Clonal exposure versus label sharing. 90.2 % of held-out cells have their exact fate vector somewhere in training, but only 47.7 % are exposed; the remaining 42.4 % match the label of an unrelated clone, which no model can use to identify them. **c**, Retrievability with 95% intervals from a clone bootstrap over 2,000 draws, which resamples query clones rather than exposed cells; counts are shown beside each point and dashed segments mark the chance level. The three intervals do not overlap. **d**, The ordering CellTag LARRY PC9 across 96 combinations of principal-component dimensionality, distance, neighborhood size, hit rule and fold design, grouped by hit rule. CellTag is the most retrievable system in all 96, and the full ordering holds in 64 of 96.

This capacity ceiling reflects ancestral lineage rather than ambient label sharing: in LARRY, 90.2 % of held-out cells share a fate vector with training cells, but for 42.4 % this occurs within unrelated clones that independently converged on that fate (Fig. 2b). The oracle establishes channel capacity when sister identities are known; actual models must discover these relatives in expression space.

### Retrievability in expression space predicts score inflation

A training relative enables data leakage only when an algorithm can locate it in expression space. We quantify this accessibility through nearest-neighbor sister retrieval *r*: the fraction of exposed held-out cells whose closest training profile in 50-dimensional PCA space of highly variable genes is a true clonal relative. Because *r* depends on representation and query rules, it is a property of the feature space rather than an intrinsic constant of the dataset.

Sister retrieval rates cleanly separate the three systems across distinct biological regimes of clonal memory (Fig. 1d). In CellTag direct reprogramming, clonal relatives remain tightly clustered in expression space, yielding *r* = 0.435 (44-fold over analytical chance 0.0098). In LARRY hematopoiesis, multi- lineage branching pulls sister cells apart, lowering retrieval to *r* = 0.039 (23-fold over chance 0.0017). In PC9 cancer persisters under osimertinib, drug selection decouples cellular state from ancestry (*r* = 0.0084, only 3-fold over chance 0.0029); for 99.2 % of exposed cells, the nearest training profile belongs to an unrelated lineage. Because *r* conditions on exposure, the joint fraction of held-out cells that both possess a training relative and retrieve it is the product *εr*: 0.3120 in CellTag, 0.0189 in LARRY, and 0.0079 in PC9. The two diagnostics rank the systems in divergent orders: exposure places PC9 first, then CellTag, then LARRY, whereas retrievability places CellTag first, then LARRY, then PC9.

To test whether retrievability predicts real benchmark inflation, we evaluated six supervised architectures across all three systems under identical preprocessing: an uninformative prior classifier, uniform *k*-NN, tuned *L*_2_ logistic regression (hereafter the tuned linear model), RBF-Nyström kernel logistic regression, gradient-boosted decision trees, and a tuned multilayer perceptron (Table 1). Holding cells, features, and tuning fixed, moving from clone-grouped to random splitting lowers held-out linear loss by 0.102 nats in CellTag, 0.011 nats in LARRY, and 0.0032 nats in PC9 (evaluation gap Δ*L*; Table 1)—a 32-fold range under one protocol. Realized score inflation strictly matches the retrievability ranking (CellTag > LARRY > PC9) across all six models on point estimates, whereas exposure would have ranked PC9 first. Resampling clones over 2,000 draws separates the CellTag gap from zero for every model of the cell-level-holdout ladder (the group-aware multilayer perceptron gap was not bootstrapped; Methods); in LARRY and PC9, the linear interval spans zero, whereas LARRY’s kernel, boosted-tree, and *k*-NN models exclude zero. Normalizing each gap to clone-grouped headroom above empirical label entropy (0.1876 in CellTag; Table 1) yields 14.0 % in CellTag, 1.5 % in LARRY, and 1.1 % in PC9 (12.6-fold range), reinforcing the retrievability ordering.

To verify that this evaluation gap reflects transcriptomic retrieval rather than fold label imbalance, we decomposed total inflation into baseline and feature-dependent components. The feature-free prior classifier retrieves nothing, yet its loss shifts between splits by 0.0195 nats in CellTag, 0.0021 in LARRY, and 0.0010 in PC9 from fold label composition. This baseline accounts for 19.0 % of the tuned linear gap in CellTag, 19.6 % in LARRY, and 30.5 % in PC9. Subtracting it isolates the feature-dependent gap (0.0830 nats in CellTag versus 0.0085 in LARRY and 0.0022 in PC9; Table 1). The ordering survives this subtraction, so the retrieval account rests on the five models that read expression.

Score inflation tracks retrievability because classifiers construct decision boundaries from local geometry: an immediate neighborhood probe captures the leak, whereas diffuse background distributions dilute it. In contrast, global metrics fail to predict vulnerability. Normalized rank-biserial distance separation, *κ* = 2 AUROC − 1, scores LARRY (*κ* = 0.309) above CellTag (*κ* = 0.287), inverting the pair whose empirical gaps differ nearly tenfold (0.102 versus 0.011 nats); only PC9 (*κ* = 0.048) matches the empirical order. Global metrics average over distant pairs, whereas nearest-neighbor retrieval *r* measures local aperture directly.

Clone bootstrap resampling confirms statistical separation among the three systems (Fig. 2c). Resampling query clones over 2,000 draws yields mutually disjoint 95% intervals: 0.3350–0.5190 in CellTag (813 retrievals across 1,871 exposed cells from 392 clones), 0.0217–0.0585 in LARRY (22 events across 564 cells from 217 clones), and 0.0055–0.0116 in PC9 (26 events across 3,093 cells from 487 clones). Heavy-tailed clone sizes widen CellTag’s interval 4.1-fold over naive estimates, while chance-adjusted retrievability, (*r* − *r*_chance_)/(1 − *r*_chance_), preserves strict separation: 0.429 in CellTag, 0.037 in LARRY, and 0.005 in PC9. Across a parameter sweep of 96 configurations varying PCA dimensionality (10 to 100), distance metrics (Euclidean, cosine), neighborhood sizes (*k* ∈ {1, 5, 10}), consensus hit rules, and fold schemes, CellTag achieves highest retrievability in all 96 conditions, and the full hierarchy holds in 64 configurations (Fig. 2d).

This hierarchy remains consistent under depth-matched refitting and clone-level reweighting. Matching sampling depth by refitting all six models on the 840 CellTag cells in 304 clones with ≥ 3 mature descendants (matching LARRY and PC9) yields a tuned linear gap of 0.0568 nats (retrievability 0.344, label entropy 0.526), exceeding LARRY’s largest gap (0.0314 nats, kernel regression) and PC9’s (0.0032 nats). Reweighting each clone equally yields tuned linear gaps of 0.0062 in CellTag, 0.0029 in LARRY, and 0.0011 in PC9, preserving the hierarchy on an independent clone scale. On the cell-weighted scale, excluding the 11 largest clones (636 cells) yields a CellTag gap of 0.0042 nats without refitting, while the depth-matched refit above confirms the hierarchy with fully refitted models.

The PC9 cohort establishes the methodological distinction between leakage opportunity and realized benchmark inflation. Despite highest exposure (*ε* = 0.943, mean 5.04 cells/clone, up to 39), PC9 inflates least (0.0032 nats). Under drug selection, persister transcriptomes decouple from lineage ancestry (*r* = 0.0084, 3-fold over chance 0.0029) [39, 40], leaving 99.2 % of exposed cells adjacent to unrelated profiles. Exposure establishes whether a leakage channel exists; retrievability bounds what can leak through it. Methodologically, validating generalization to unseen clones requires grouped holdout, and a small measured gap provides empirical evidence about the cohort rather than justification for cell-level splitting.

### Two arms of a single study diverge 25.1-fold under an identical protocol

Evaluating contrasting biological processes within a single study isolates clonal architecture from technical confounders. In Jindal et al. [37], two arms share an identical protocol: mouse LSK hematopoietic progenitors profiled at day 2.5 with fate recorded at day 5 (75.6 % published RNA-only accuracy), and fibroblasts undergoing direct reprogramming to induced endoderm progenitors at day 3 (fate at days 12 and 21). Both arms specified five-fold stratified cross-validation with five repeats. Holding cells, features, and estimators fixed, we paired this with clone grouping.

Pre-flight diagnostics reveal opposite extremes of vulnerability. LSK hematopoiesis (1,520 cells, 1,198 clones, 6 classes) exhibits modest exposure (*ε* = 0.325) and low retrievability (*r* = 0.021; chance 0.0010; 21-fold enrichment). Direct reprogramming (3,143 cells, 1,290 clones, 3 classes) exhibits high exposure (*ε* = 0.716) and high retrievability (*r* = 0.409; chance 0.0085; 48-fold enrichment). Clonal architecture explains this contrast: LSK progenitors divide briefly yielding small, mostly singleton clones, whereas reprogramming generates heavy-tailed expansions clustered in expression space. The reprogramming arm is drawn from the released clone table and kept separate from the 2,606-cell cohort.

Three independent anchors verify reproduction fidelity before evaluating the split. First, a highly variable gene model reproduces published accuracy within 3.1 percentage points (0.7254 versus 0.7564). Second, a transcription-factor model matches published performance (0.6511 versus 0.638), spanning an 11.8 percentage-point range across representations. Third, our pipeline recovers the authors’ 1,422 state–fate clones exactly without tuning, confirming identical cohort definition.

Evaluating both arms under clone-grouped holdout confirms that benchmark inflation tracks retrievability. In hematopoiesis, published performance transfers to held-out clones: accuracy is 0.7254 under random splitting versus 0.7194 grouped (gap 0.006; clone bootstrap 95% CI: 0.0015–0.0106 over 2,000 draws). In reprogramming, the gap widens to 0.151 fold-averaged (0.7092 random versus 0.5578 grouped) and 0.1615 pooled per cell (95% CI: 0.0872–0.2356). Identical code yields a 25.1-fold divergence in evaluation gap, anticipated before model training by a 19.4-fold difference in retrievability *r*.

Clone-label permutation shows that instance lookup alone can produce inflation of this size in the reprogramming arm. Permuting fate labels across intact clones breaks biological associations while preserving clonal geometry: random splitting retains 0.5511 accuracy over 30 replicates against a permuted base rate of 0.4657, whereas clone-grouped validation falls to chance (0.4386). The resulting permuted gap is 0.1125 ± 0.0448 (2.5th–97.5th percentile 0.0309–0.1763; positive in every replicate, minimum 0.0298). In hematopoiesis, permutation reduces both splits to chance base rates (0.5989 and 0.5992 versus 0.6000). Across 12 configurations varying features and estimators (8 in hematopoiesis, 4 in reprogramming), disjoint distributions (−0.0008 to 0.0071 versus 0.1490 to 0.1708) confirm that this divergence is robust.

### Donor-level scoring reveals whether evaluation gaps alter clinical conclusions

We next examine a clinical cohort with patient-level diagnostic labels. Because each donor contributes hundreds of cells, exposure saturates (*ε* = 1 across folds) under cell-level random cross-validation. In translational medicine, clinical utility requires generalization to unseen patients rather than profiled donors. An open leakage channel allows models to exploit patient identity shortcuts; the scoring unit therefore dictates benchmark fidelity. We evaluate models at both patient and cell levels to establish whether evaluation gaps alter diagnostic conclusions.

Postmortem human brain microglia from the Seattle Alzheimer’s Disease Brain Cell Atlas (SEA- AD) [41] (85 donors: 40 dementia, 45 control; 34,000 cells, 10 folds repeated 5 times) exhibit an open leakage channel (*ε* = 1.000, *r* = 0.439; 37-fold over chance 0.0117). At the patient level, random splitting classifies 73.8 of 85 donors correctly (AUROC 0.9641), whereas donor-grouped holdout correctly classifies 59.2 (AUROC 0.7553)—an evaluation gap of 0.209 in AUROC (Fig. 3a,b; donor bootstrap gap 0.1878, 95% CI: 0.1198–0.2649 over 2,000 draws). Scored per cell, models report AUROCs of 0.8315 and 0.6741 (gap 0.157; Fig. 3c; bootstrap 95% CI: 0.118–0.180). Cell-level scoring understates the clinical gap (0.157 versus 0.209), showing that benchmark fidelity depends on the evaluation unit.

**Figure 3:**
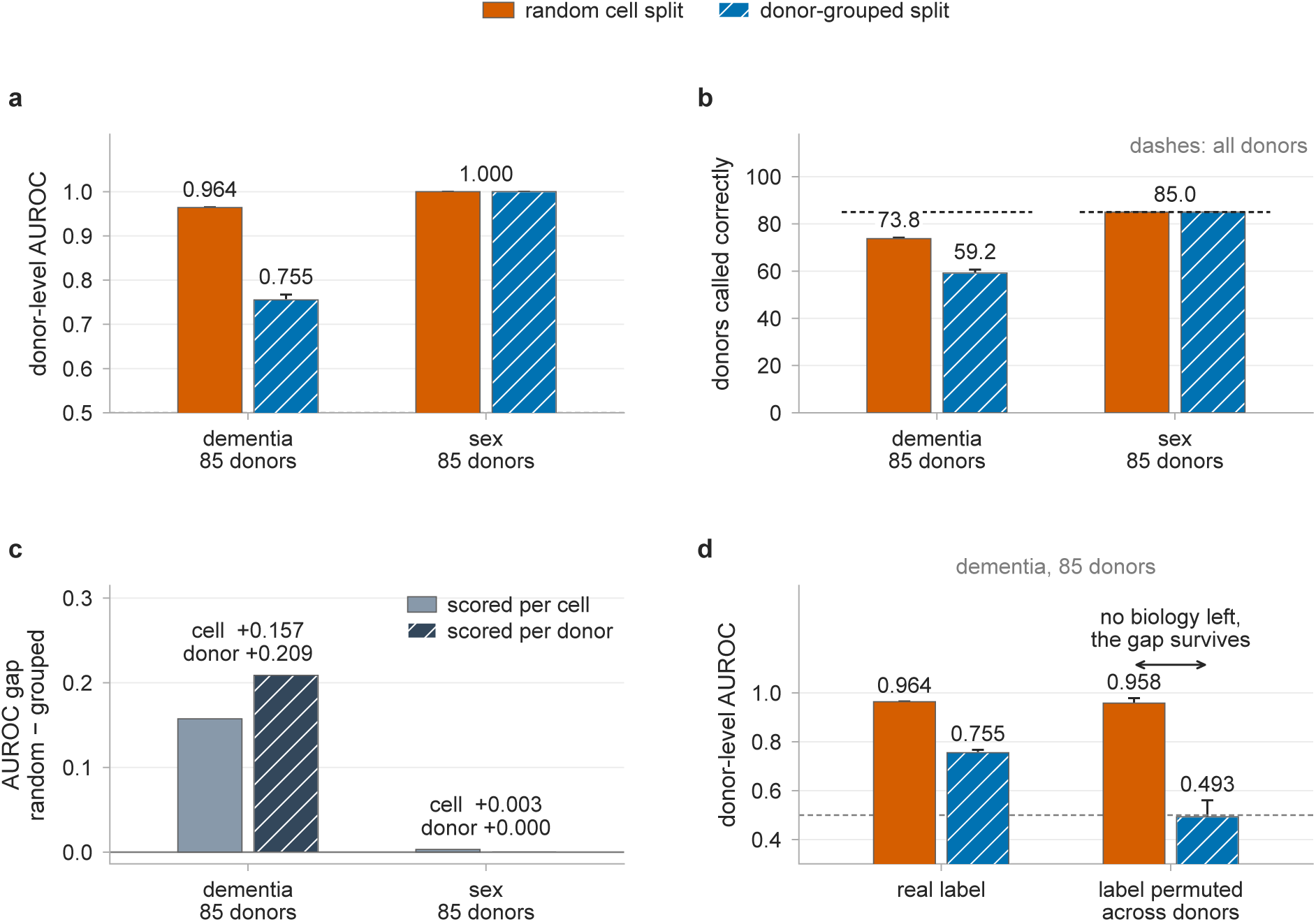
Donor-owned labels, scored at the unit that owns them in SEA-AD microglia. Cells assigned at random (orange, solid) against donors held out whole (blue, hatched) across 85 donors; everything else (cells, features, estimator, fold count) is identical between arms. Error bars in **a** and **b** are the standard deviation over repeats (5 for dementia, 3 for sex); in **d** they are the standard deviation over the 20 label permutations. Panel **c** plots a difference and is direct-labeled instead. **a**, Donor-level AUROC, computed on each donor’s mean predicted probability over that donor’s held-out cells. Dementia falls from 0.9641 to 0.7553, whereas donor sex (positive control on the identical donors) remains at 1.0000 (gap 0.000). **b**, Donors called correctly, with the cohort size marked by a dashed line: 73.8 against 59.2 of 85 in dementia, and 85.0 of 85 under both arms for sex. **c**, The same gap scored two ways. Scoring per cell understates the dementia gap (0.157 against 0.209): the unit at which a metric is computed changes what it reports. **d**, Dementia under a donor-label permutation (20 replicates), which destroys the biology while preserving each donor’s cells as a block. The random split still reaches a donor AUROC of 0.958 and the grouped split falls to 0.493. Permuting labels across cells instead sends both arms to cell-level AUROC 0.501 and 0.501 (not plotted; donor-level AUROC is undefined once a donor no longer owns its label).

Permutation controls show that donor identity alone can generate an evaluation gap of this size. Shuffling disease labels across intact donor blocks preserves data geometry while breaking disease associations: random splitting still attains donor AUROC 0.958 (75.3 donors correct), whereas donor-grouped validation collapses to chance (0.493; Fig. 3d; 20 replicates). At the cell level, permuted arms yield a gap of 0.249 (0.748 versus 0.498), exceeding the unpermuted gap of 0.157. Conversely, shuffling labels across individual cells uncouples donor identity, driving both splits to chance (0.501 and 0.501).

Holding donors out preserves full performance when biological signal is cell-intrinsic. Because sex chromosome genes (such as *XIST* in females) mark cells cell-autonomously, donor sex serves as a positive control on the identical 85 SEA-AD donors (49 female, 36 male; 34,000 cells, 2,000 features, *ε* = 1.000, *r* = 0.439; Table 2). Sex classification transfers across donor holdout without loss: donor AUROC is 1.0000 random versus 1.0000 grouped (85.0 of 85 donors correct random versus 85.0 grouped; gap exactly 0.000; Fig. 3a,b; cell-level gap 0.0030, from 0.9958 to 0.9927). Shifting the target from sex to dementia on this identical channel expands the donor gap from 0.000 to 0.209, showing that channel geometry bounds the gap while target biology determines how much passes through it.

**Table 2:** The same shape in two regimes. Exposure and retrievability are computed with no label and no model, on the random arm; the grouped arm has 0 by construction. The two clone-owned rows come from one published study with one described protocol; the study’s published RNA-only accuracy, 75.6 %, belongs to the hematopoiesis row. ^†^Donor sex serves as a cell-intrinsic positive control on the identical 34,000 cells and donor structure as dementia; its retrievability is the same measurement as the dementia row.

| Task and cohort | $\epsilon$ | $r$ | Impact of grouped holdout |
| --- | --- | --- | --- |
| *Label owned by a clone: two arms of one study, one described protocol* |  |  |  |
| Reprogramming, CellTag-multi | 0.716 | 0.409 | Accuracy 0.7092 → 0.5578; gap +0.151 per fold, +0.1615 pooled per cell (bootstrap 0.0872–0.2356 on the pooled form) |
| Hematopoiesis, CellTag-multi | 0.325 | 0.021 | Accuracy 0.7254 → 0.7194; gap +0.006, bootstrap 0.0015–0.0106 |
| *Label owned by a donor* |  |  |  |
| Dementia, SEA-AD, 85 brains | 1.000 | 0.439 | Donor AUROC 0.9641 → 0.7553; 73.8 → 59.2 donors called correctly |
| Sex, SEA-AD, 85 brains | 1.000 | 0.439<sup>†</sup> | Donor AUROC 1.0000 → 1.0000; gap exactly 0.000 |

Altering label ownership reinforces this principle. For cell-type annotations owned by individual cells (the 8 SEA-AD microglia/PVM supertypes in this cohort), overall accuracy between splits agrees within 0.0078 over 5 repeats. Balanced accuracy shifts by 0.042 (0.699 to 0.658), with seven of eight states shifting by at most 0.025; the residual change is driven by a single rare 156-cell state whose recall falls from 0.474 to 0.186 due to training donor exclusion rather than donor ownership. Where labels belong to individual cells rather than donors, grouped validation preserves benchmark performance, confirming that evaluation gaps stem from label ownership rather than grouping per se.

### Representation capacity bounds realized leakage

Data leakage requires both an open channel in data geometry and a feature representation sufficiently expressive to exploit it. We hypothesized that low-dimensional summaries bottleneck the channel, preventing shortcut memorization even when data exposure is high. In CellRank’s benchmark on mouse embryonic fibroblast reprogramming [7, 45], binary fate is predicted from a scalar velocity-derived fate probability. Across 3,049 labeled cells with ground-truth CellTag clone fates, we evaluated three nested representations—scalar probability, 50 principal components, and 2,000 genes—measuring out-of-fold AUROC gaps across 25 resampled draws.

On a scalar input, data leakage is strictly bounded by representation capacity. Because one-dimensional logistic regression is a monotonic transformation, out-of-fold AUROC analytically equals the raw input AUROC oriented by coefficient sign; classifier and oriented feature AUROCs agree in 2,367 of 2,369 fits to within 1.67 × 10^−16^ (2 zero-coefficient exceptions). For this ranking metric, the model transmits at most a single bit—boundary orientation—and cannot encode instance identities. The AUROC evaluation gap remains negligible across all three clone definitions (0.018 fine, 0.026 coarse, and -0.014 connected-component; Fig. 4a), whereas in 50-dimensional PCA space exposure and retrievability are high (*ε* = 0.940, *r* = 0.443, product *εr* = 0.416, day-21 fine clones; Fig. 4b). Under label permutation, this one-bit channel leaves a minimal residual AUROC gap of only 0.072 (0.579 random versus 0.507 grouped; Fig. 4c).

**Figure 4:**
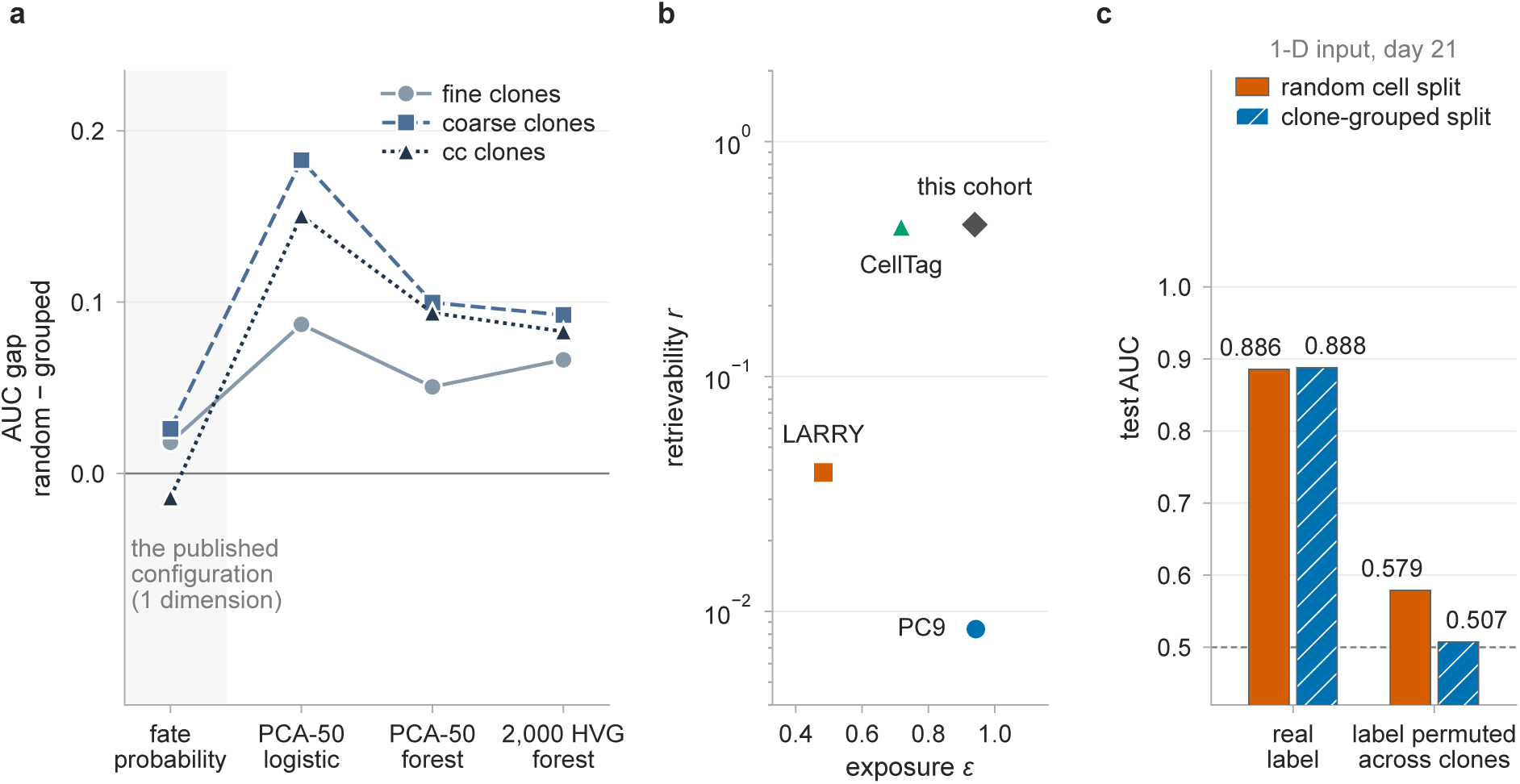
Representation capacity bounds transmission through the leakage channel. Binary fate prediction across 3,049 lineage-labeled cells in mouse embryonic fibroblast direct reprogramming [7, 45], comparing feature representations on identical cells and partitions. **a**, Measured AUC gap between a random split and a clone-grouped one over 25 draws, averaged across three timepoints, holding the cells, the labels and both fold assignments fixed and changing only the input. With the published one-dimensional input (shaded) the gap is near zero under all three clone definitions; with 50 principal components and the same logistic head it reaches 0.087 to 0.183, and neither a forest on those components nor a forest on 2,000 genes exceeds it. **b**, The channel is open: measured in the same 50-dimensional PCA reference space, the product *εr* = 0 416 places this cohort above every barcoded system analyzed here, CellTag included. **c**, The channel is measurable even with a one-dimensional input. Permuting the label across clones removes all real signal, and the random split still scores AUROC 0.579 against 0.507, the residual a monotone scalar can transmit when only its boundary orientation is fitted.

Expanding feature capacity immediately opens the channel. Providing the same logistic classifier with 50 principal components produces large AUROC gaps (0.087 fine, 0.183 coarse, and 0.150 connected-component), supplying capacity to isolate individual cells. Increasing model flexibility yields comparable inflation: random forests on 50 components (AUROC gap 0.051) and on 2,000 genes (0.066; Fig. 4a) extract no more leakage than linear models on 50 components. Moving from fate probability to principal components provides expression coordinates where clone identity can be read; greater model flexibility cannot extract information absent from the input representation.

### Grouped splitting realigns precision with the units that vary

Holding out whole groups closes the leakage channel, but changes the statistical unit of error estimation. Holding clones out increases fold-to-fold loss standard deviation by 1.30-fold in LARRY, 2.08-fold in PC9, and 6.56-fold in CellTag (rising from 0.021 to 0.135 nats in CellTag), driven by heavy-tailed clone sizes (109 cells in CellTag versus 39 in PC9 and 7 in LARRY; Fig. 1b). Consequently, direct re-programming is simultaneously where grouping is most informative and where fold variance is highest.

Conversely, cell-level random splitting underestimates standard errors by 1.39-fold in LARRY, 3.15-fold in PC9, and 6.59-fold in CellTag relative to cluster-robust sandwich estimators. The effective sample size for these loss estimates is 608 independent units for LARRY’s 1,167 cells, 331 for PC9’s 3,280 cells, and 60 for CellTag’s 2,606 cells. This quantity describes the estimate, not the dataset (CellTag has 1,066 clones), reflecting the degrees of freedom an error bar on this loss can legitimately claim. Grouped validation aligns statistical precision with the independent biological units that actually vary.

When biological groups are few, grouping also sets fold class composition. In SEA-AD donor sex (85 donors), 1 of 30 grouped folds has a single class, requiring pooled reporting. Under leave-one-group-out validation in CellRank reprogramming, only 3 of 186 clone holdouts, pooled over three outcome timepoints and three clone definitions, admit an AUROC, and none of the 114 holdouts under the finest definition does. This reflects the limitation of per-fold ranking metrics under small test cohorts; pooling out-of-fold predictions or employing proper scoring rules resolves this issue directly.

### Pre-flight profiling with leakcheck

To operationalize these geometric diagnostics prior to model fitting, we implement closed-form exposure *ε* and nearest-neighbor retrieval *r* in `leakcheck`, an open-source Python tool. Taking a grouping vector (such as clones, donors, batches, or wells) and a PCA matrix, `leakcheck` computes (*ε*, *r*) coordinates, analytical chance expectations, and group-size distributions in seconds without requiring model training or label access. This diagnostic enables practitioners to profile evaluation vulnerability pre-flight and report standardized (*ε*, *r*) coordinates in benchmark evaluations across studies.

## Discussion

Random cross-validation on grouped single-cell data creates silent leakage: whenever test cells share clonal lineages, patient donors, or outcome-linked experimental batches with the training partition, predictive models can achieve inflated benchmark performance by memorizing training relatives instead of learning transferable biology. Here we show that this leakage vulnerability is not uniform, but bounded by two geometric properties of the data that are computable before training any outcome model: exposure (*ε*), which determines whether a random split opens a leakage channel, and retrievability (*r*), which determines how much can leak through it in expression space. We implement these pre-flight diagnostics in `leakcheck`, which returns both coordinates along with chance baselines and group distributions in seconds from feature matrices and group keys alone.

Across empirical benchmarks, when relatives cluster tightly in gene expression space (high retrievability *r*), random splitting allows models to exploit nearest-neighbor shortcuts, producing marked performance inflation. Conversely, when cell states decouple from lineage ancestry, nearest-neighbor queries return unrelated cells, leaving little for a random split to exploit. Furthermore, even when a leakage channel is open, models generalize without loss when target signals reflect cell-intrinsic biology rather than patient identity shortcuts. Scoring at the label-owning unit under both splits resolves whether high performance reflects generalizable biology or shortcut learning. Exposure and retrievability bound this channel directly from data geometry, before fitting an outcome model.

Validation design must therefore align with the scientific claim: evaluating whether a model generalizes to unseen clones, patients, or experimental batches whose composition is tied to outcome requires holding those units out. A small measured gap is then evidence about the cohort in hand, not a justification for treating related cells as independent. Where groups are held out, grouped cross-validation realigns statistical precision with the independent biological units that actually vary, preventing pseudoreplication while managing fold variance.

For practitioners, pre-flight profiling with `leakcheck` assesses leakage risk before model training. These coordinates describe data geometry under one representation, neighbor rule, and fold design. Extending this ordering to calibrated gap predictions will benefit from prospective validation across external cohorts, deeper target-support thresholds, and refit-inclusive group bootstraps. Reporting continuous (*ε*, *r*) coordinates documents evaluation vulnerability alongside benchmark scores.

## Methods

### Three consequences of group structure, kept separate

Group structure influences a cell-level split in three ways that are distinguished throughout. *Leakage* is the presence in the training partition of target information about a specific held-out cell (here, the label of a clonal sister or same-donor cell) that an algorithm can retrieve and exploit, artificially inflating test scores. *Pseudoreplication* is the treatment of correlated cells as independent observations; for a fixed cell-weighted estimand, it leaves the point estimate untouched and understates its standard error, quantified here via design effects and effective sample sizes. *Distribution shift* is the change in feature distribution that holding out whole groups additionally causes, which can move scores in either direction independently of lookup. Label permutations show what the first can produce on its own by destroying biological associations while preserving data geometry; cluster-robust standard errors address pseudoreplication directly.

### Datasets and experimental provenance

All single-cell transcriptomic datasets and clonal barcode annotations were obtained from published repositories and staged under cryptographic SHA-256 verification. For LARRY [36], *in vitro* day-2 mouse hematopoietic stem and progenitor cells were profiled to predict mature day-4/6 lineage fate across 6 classes (GEO GSM4185642). Cells with exactly one barcode and ≥ 3 mature clonal descendants were retained (1,167 cells, 785 clones). For CellTag-multi [37], day-3 mouse embryonic fibroblasts undergoing direct lineage conversion to induced endoderm progenitors were used to predict terminal differentiation into 3 outcomes defined by day-12/21 descendants (2,606 cells, 1,066 clones). For PC9 [38], day-3 human EGFR-mutant lung adenocarcinoma cells treated with osimertinib were used to predict each lineage’s day-14 persister outcome; lineages with at least 3 scored descendants were retained (3,280 cells, 651 lineages). That target is a scalar in [0, 1], representing the fraction of a lineage’s day-14 descendants recovered in the cycling gate, with the remainder non-cycling persisters; the two-term cross-entropy is symmetric in which fraction is named.

The three cohorts differ in how many mature descendants support each clone’s fate vector. LARRY and PC9 retain only lineages with at least three, giving a median target support of 6 descendants per cell in LARRY and 6 in PC9. In contrast, CellTag includes all descendants: median target support is 1 descendant, and 1,414 of its 2,606 cells (54.3 %), from 609 of 1,066 clones, have a fate vector estimated from a single day-12/21 descendant, giving zero empirical entropy by construction. Because a random split can reproduce a clone’s sampling realization from a training sister while a grouped split cannot, all six models were refitted on the 840 CellTag cells in 304 clones that meet the three-descendant criterion. Fold assignment, highly variable gene selection, scaling, PCA, and hyperparameter search were repeated inside this subset. Exposure there is 0.799, retrievability 0.344 (against 0.435 on the full cohort), mean label entropy 0.526, and the tuned linear gap 0.0568 nats; the ordering across the three systems holds for every model. The label entropy used to normalize Table 1 is the entropy of these empirical fate vectors, providing a floor for the observed target rather than a bound on biological uncertainty.

Held-out loss is out-of-fold soft cross-entropy in nats, averaged over cells, so that larger clones contribute more,

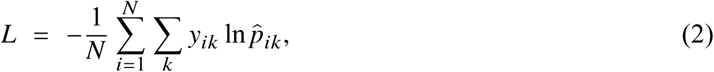

with the two-term binary form for PC9; predicted probabilities are smoothed by empirical prior blending *p̂* = (1 −*ε*_prior_)*P* +*ε*_prior_ prior, *ε*_prior_ = 10^−3^, and floored at 10^−12^. The evaluation gap, or score optimism, is Δ*L* = *L*_grouped_ − *L*_random_, with *L*_grouped_ the clone-grouped and *L*_random_ the cell-level random loss on the same cells. Fold assignment is made once, at the clone level, by largest-first load balancing on cell count; the random arm permutes cells into five folds of the same sizes, so the two arms differ in the assignment and not in the fold geometry. Because fold sizes are fixed rather than drawn per cell, the exact expectation of exposure is hypergeometric rather than the closed form quoted in the text. For a test fold of size *n_k_* drawn from *N* cells, a query cell from a clone of size *m* has no training relative with probability

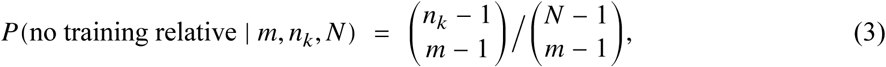

and exposure is one minus the cell-weighted average of that term over clones and folds. The leakage dose *λ* of Fig. 1c interpolates between the two arms by dissolving a fraction *λ* of the multi-cell clones into single-cell units before fold assignment, in nested subsets at a fixed seed, so that a cell moves only when its own clone dissolves; the curve is the mean over three seeds.

### The six models and hyperparameter tuning

We evaluated a fixed panel of 6 models across all three systems: the prior classifier predicting the training-fold base rate, uniform-weighted *k*-NN in 50-dimensional PCA space (*k* ∈ {5, 15, 50}), tuned multinomial *L*_2_ logistic regression (*C*_reg_ ∈ {10^−5^, …, 10^3^} across 9 logarithmic values), RBF kernel logistic regression via scikit-learn’s `Nystroem` approximation (min(300, *N*_train_) com-ponents, *γ* ∈ {0.003, 0.01, 0.03}, *C*_reg_ ∈ {0.1, 1, 10}), histogram-based gradient-boosted trees over (max_leaf_nodes, learning_rate) ∈ {(4, 0.1), (8, 0.1), (16, 0.03)}, and a multilayer perceptron (MLP) with two equal-width hidden layers (ℎ ∈ {32, 128}), ReLU activations, dropout *p* = 0, Adam, weight decay ∈ {10^−3^, 10^−2^}, and a 20 % validation holdout for early stopping drawn over whole clones. All models use standard scikit-learn estimators and a PyTorch MLP [33, 43]. Highly variable gene selection, scaling, and PCA were fitted within the training partition of each fold; hyperparameters were tuned in an internal 3-fold loop grouped by clone. With the holdout drawn over cells instead, the CellTag gap is 0.2803 nats, LARRY’s 0.0168 and PC9’s 0.0044, leaving the ordering unchanged. The clone-bootstrap intervals in Table 1 and the text were computed on the cell-level-holdout runs; the group-aware re-run stored per-seed means only and was not bootstrapped. The tuned linear model is group-aware at every step, providing a rigorously tuned baseline.

### Within-clone transcriptomic similarity and PCA space

The top 2,000 highly variable genes were selected by dispersion: for the outcome models on the raw counts of each training fold (on the log-normalized cache for PC9), and for the retrievability probe on log-normalized values over the whole cohort, so the probe’s feature space is close to, but not identical with, the one each fitted model receives. Raw unique molecular identifier (UMI) counts were normalized to a constant library size (10^4^ per cell), transformed by ln(1 + *x*), scaled to zero mean and unit variance, and projected onto 50 principal components [46]. Exposure is the fraction of held-out units with at least one same-group member in the training partition, pooled over folds. For every held-out cell with at least one clonal sister in the training partition (an exposed cell), we queried its single nearest training neighbor by Euclidean distance in that 50-dimensional space. The retrieval rate *r* is the fraction of exposed cells whose nearest training neighbor is a true sister. Chance retrievability is the analytical probability that a uniformly drawn training cell shares the query cell’s clone, *r*_chance_ = ∑*_c_ N*_test,*c*_(*N*_train,*c*_/*N*_train_)/*N*_exposed_. Uncertainty on *r* is a percentile bootstrap over the clones that own the query cells, resampling whole clones with replacement over 2,000 draws and recomputing the rate on the resampled set. Chance-adjusted retrievability is (*r* − *r*_chance_)/(1 − *r*_chance_), the fraction of the achievable margin above chance that the retrieval realizes.

In the workflow `leakcheck` implements, the PCA handed to the probe is fitted on the full dataset before fold assembly as a target-blind structural diagnostic. Sensitivity checks across 96 combinations of PCA dimensionality (10 to 100), distance metrics (Euclidean, cosine), neighborhood sizes (*k* ∈ {1, 5, 10}), hit rules, and fold schemes confirmed stability without fitting outcome models. Transgene-derived features are present in two matrices (CellTag.UTR and GFP.CDS in CellTag; H2B-mCherry in PC9) and are selected among the 2,000 highly variable genes in every fold. Excluding them leaves the tuned linear gap at 0.1029 nats in CellTag (against 0.1024) and 0.0032 in PC9 (against 0.0032), and the CellTag *k*-NN gap at 0.2414 (against 0.2409); LARRY carries no such feature. The probe executes in 0.011 seconds on 3,280 cells (median of 25 warm calls on one AMD Ryzen 9 9950X 16-Core Processor; 0.32 seconds cold start), remaining under one second through 60,000 cells.

### The sister-lookup oracle

The oracle fits nothing: for each held-out cell it predicts the fate vector of a clonal sister in the training partition when available, returning the cell’s own label with cross-entropy *H*(*y_i_*), and the training-fold base rate otherwise. Evaluated under clone-grouped splitting, the sister branch is never available. Both numbers are means over the same 1,167 cells and the same 5 seeds, using the fold assignments of the two endpoints. Because the oracle redraws its random split at each seed, the exposure it sees (0.477) is a mean over those seeds, aligning with the canonical 0.483 measured on the primary fold assignment. The oracle bounds the information the leakage channel supplies to a predictor that uses nothing else. It is not an upper bound on an arbitrary model: an expression-based predictor may exceed the oracle without exploiting any leakage at all.

### Statistical estimation and variance analysis

Confidence intervals for score optimism are percentile cluster bootstraps over clone identifiers on stored out-of-fold per-cell losses, reflecting held-out cell uncertainty conditional on fitted predictions rather than split-to-split retraining variation (reported as seed-to-seed and fold-to-fold spread). Cluster-robust standard errors use the Huber–White sandwich estimator clustered by clone ID. Effective sample sizes use the design effect measured directly from the two standard errors on the same held-out losses, deff = (SE_cluster_/SE_naive_)^2^ and *N*_eff_ = *N*/deff, accommodating heavy-tailed clone-size distributions directly.

### The two arms of the CellTag-multi study

For hematopoiesis [37], mouse LSK progenitors were profiled at day 2.5 with clone-level fate read at day 5, restricted to the authors’ released barcode list (1,520 cells in 1,198 clones over 6 collapsed fate classes). For reprogramming, fibroblasts under forced conversion were profiled at day 3 with clone-level fate from the released clone table (3,143 cells in 1,290 clones over 3 classes). Primary features are scanpy’s analytic Pearson residuals over 3,000 highly variable genes; accuracy is within 3.1 points of published performance, and the gap remains small across all 8 configurations.

Hyperparameters were frozen from the published random-forest grid search: arm A reproduces the published cell-level splitter (`RepeatedStratifiedKFold(n_splits=5, n_repeats=5, random_state=0)`), while arm B uses `StratifiedGroupKFold(n_splits=5, shuffle=True)` with `groups = clone` over five random states (25 fits per arm; inner re-tuning yields 0.1490 gap). The clone bootstrap resamples clones with replacement over 2,000 draws (centering at 0.1615 versus fold-averaged 0.1514). Clone-label permutations shuffle fate labels across intact clones over 30 replicates in reprogramming and 3 in hematopoiesis.

### Donor cohorts

We analyzed SEA-AD microglia [41] from dorsolateral prefrontal cortex and middle temporal gyrus (dementia versus control). Cells per donor were equalized by subsampling (400 in brain, seed 0) allocated proportionally across libraries; donors below the floor were excluded, leaving 85 of 89 brain donors (34,000 of 82,486 cells). Counts were normalized to 10^4^ per cell, ln(1 + *x*)-transformed, and filtered to the top 2,000 highly variable genes by `seurat_v3` identically between arms. Gene selection is transductive: identical between arms without labels or fold assignments, it cannot generate the gap between them, as held-out donor expression enters both arms identically. A sensitivity run selecting 2,000 genes strictly inside each outer training fold over 5 repeats confirmed donor-level gaps of 0.2083 in dementia (versus 0.209 shared) and exactly 0.000 for sex. The estimator is *L*_2_ logistic regression with balanced class weights and fold-internal scaling, evaluated across 10-fold random and donor-grouped splits over 5 disease repeats and 3 sex repeats (1 of 30 grouped folds contained a single class, evaluated via pooled metrics). Cell-type controls evaluated microglial annotations over 5 repeats.

Donor-level metrics average predicted probabilities across each donor’s held-out cells, counting a donor correct when mean probability exceeds 0.5. Donor bootstraps resampled donors over 2,000 draws on out-of-fold predictions. Evaluating a single repeat’s pooled predictions captures donor sampling rather than fold variation (both point estimates and across-repeat means are given). Donor-label permutations shuffled labels across intact donors over 20 replicates. Replacing ten-fold grouping with leave-one-donor-out yields a cell-level gap of 0.163 in dementia (against 0.157 at ten folds). Within-brain-region neighbor queries yield *r* = 0.477 (versus 0.439 cohort-wide), confirming that retrievability measures donor identity rather than brain region.

### The one-dimensional configuration

The CellRank reprogramming cohort profiles mouse embryonic fibroblasts undergoing direct conversion to induced endoderm progenitors tagged with CellTag barcodes [7, 45]. We analyzed the distributed 48k subset restricted to 3,049 cells with binary outcome labels (successful reprogramming versus dead end) on the published 60/40 train–test partition (`test_size = 0.4`) with scalar velocity-derived fate inputs [47]. Three clone definitions were constructed from object CellTag columns (fine, coarse, connected components). Across 25 invariant draws, out-of-fold AUROC matches raw feature AUROC oriented by coefficient sign within 1.67 × 10^−16^ in 2,367 of 2,369 fits (2 zero-coefficient exceptions).

### Software defaults and the grouped-by-construction literature

Standard cross-validation partitions matrix rows independently unless group-aware splitters are explicitly invoked with grouping keys, even when metadata record donor or clonal hierarchies. In contrast, multiple-instance learning architectures treat donors as intact samples by construction, making cell-level splitting structurally unavailable [23, 24, 25, 26, 27, 28, 29, 30, 31] and establishing precedents for donor-grouped evaluation.

## Data availability

All analyzed datasets are public: GEO GSE140802 (LARRY, GSM4185642); GSE216521 (CellTag-multi; induced-endoderm arm GSE216518, LSK hematopoiesis arm GSE216606); GSE150949 (PC9); SEA-AD microglia; and CellRank’s 48k subset of Biddy et al. (GEO GSE99915). Staged files and tables are available on request.

## Code availability

`leakcheck` is released as an open-source Python module with tests and a worked example at https://github.com/hucang0/leakcheck. Analysis pipelines, staged datasets, and result files are available on request.

## Acknowledgements

We thank the investigators who generated and shared the datasets analyzed here: LARRY hematopoiesis, CellTag-multi reprogramming, PC9 persisters, SEA-AD microglia, and the Biddy et al. cohort. Research reported in this publication was supported by the National Human Genome Research Institute of the National Institutes of Health under Award Number R01HG014004.

## Author contributions

H.C. conceived and designed the study, developed the theoretical framework, implemented the pipeline and tool, performed evaluations, and wrote the manuscript. S.S. contributed to conceptual development, biological interpretation, and manuscript editing. Both authors approved the final manuscript.

## Competing interests

The authors declare no competing interests.

## Notes

### Competing Interest Statement

The authors have declared no competing interest.

## References

[1] Haotian Cui, Chloe Wang, Hassaan Maan, Kuan Pang, Fengning Luo, Nan Duan, and Bo Wang. scGPT: toward building a foundation model for single-cell multi-omics using generative AI. Nature Methods, 21:1470–1480, 2024. PMID 38409223.

[2] Christina V. Theodoris, Ling Xiao, Anant Chopra, Mark D. Chaffin, Zeina R. Al Sayed, Matthew C. Hill, Helene Mantineo, Elizabeth M. Brydon, Zexian Zeng, X. Shirley Liu, and Patrick T. Ellinor. Transfer learning enables predictions in network biology. Nature, 618:616–624, 2023. PMID 37258680; PMC10949956.

[3] Minsheng Hao, Jing Gong, Xin Zeng, Chiming Liu, Yucheng Guo, Xingyi Cheng, Taifeng Wang, Jianzhu Ma, Xuegong Zhang, and Le Song. Large-scale foundation model on single-cell transcriptomics. Nature Methods, 21:1481–1491, 2024. PMID 38844628.

[4] Romain Lopez, Jeffrey Regier, Michael B. Cole, Michael I. Jordan, and Nir Yosef. Deep generative modeling for single-cell transcriptomics. Nature Methods, 15:1053–1058, 2018.

[5] Fan Yang, Wenchuan Wang, Fang Wang, Yuan Fang, Duyu Tang, Junzhou Huang, Hui Lu, and Jianhua Yao. scBERT as a large-scale pretrained deep language model for cell type annotation of single-cell RNA-seq data. Nature Machine Intelligence, 4:852–866, 2022.

[6] Shou-Wen Wang, Michael J. Herriges, Kilian Hurley, Darrell N. Kotton, and Allon M. Klein. CoSpar identifies early cell fate biases from single-cell transcriptomic and lineage information. Nature Biotechnology, 40:1066–1074, 2022. PMID 35190690.

[7] Marius Lange, Volker Bergen, Michal Klein, Manu Setty, Bernhard Reuter, Mostafa Bakhti, Heiko Lickert, Meshal Ansari, Janine Schniering, Herbert B. Schiller, Dana Pe’er, and Fabian J. Theis. CellRank for directed single-cell fate mapping. Nature Methods, 19:159–170, 2022. PMID 35027767; PMC8828480.

[8] Philipp Weiler, Marius Lange, Michal Klein, Dana Pe’er, and Fabian Theis. CellRank 2: unified fate mapping in multiview single-cell data. Nature Methods, 21:1196–1205, 2024. PMID 38871986; PMC11239496.

[9] Grace Hui Ting Yeo, Sachit D. Saksena, and David K. Gifford. Generative modeling of single-cell time series with PRESCIENT enables prediction of cell trajectories with interventions. Nature Communications, 12, 2021. PMID 34050150; PMC8163769.

[10] Mehrshad Sadria, Allen Zhang, and Gary D. Bader. Deep Lineage: Single-Cell Lineage Tracing and Fate Inference Using Deep Learning, 2024. Preprint at 10.1101/2024.04.25.591126.

[11] Sudhir Varma and Richard Simon. Bias in error estimation when using cross-validation for model selection. BMC Bioinformatics, 7, 2006. PMID 16504092; PMC1397873.

[12] David R. Roberts, Volker Bahn, Simone Ciuti, Mark S. Boyce, Jane Elith, Gurutzeta Guillera-Arroita, Severin Hauenstein, José J. Lahoz-Monfort, Boris Schröder, Wilfried Thuiller, David I. Warton, Brendan A. Wintle, Florian Hartig, and Carsten F. Dormann. Cross-validation strategies for data with temporal, spatial, hierarchical, or phylogenetic structure. Ecography, 40:913–929, 2017.

[13] Sylvain Arlot and Alain Celisse. A survey of cross-validation procedures for model selection. Statistics Surveys, 4, 2010.

[14] Gaël Varoquaux. Cross-validation failure: Small sample sizes lead to large error bars. NeuroImage, 180:68–77, 2018.

[15] Russell A. Poldrack, Grace Huckins, and Gael Varoquaux. Establishment of Best Practices for Evidence for Prediction. JAMA Psychiatry, 77:534, 2020.

[16] Stanley E Lazic. The problem of pseudoreplication in neuroscientific studies: is it affecting your analysis? BMC Neuroscience, 11, 2010.

[17] Emmeke Aarts, Matthijs Verhage, Jesse V Veenvliet, Conor V Dolan, and Sophie van der Sluis. A solution to dependency: using multilevel analysis to accommodate nested data. Nature Neuroscience, 17:491–496, 2014.

[18] Jordan W. Squair, Matthieu Gautier, Claudia Kathe, Mark A. Anderson, Nicholas D. James, Thomas H. Hutson, Rémi Hudelle, Taha Qaiser, Kaya J. E. Matson, Quentin Barraud, Ariel J. Levine, Gioele La Manno, et al. Confronting false discoveries in single-cell differential expression. Nature Communications, 12, 2021.

[19] Kip D. Zimmerman, Mark A. Espeland, and Carl D. Langefeld. A practical solution to pseudoreplication bias in single-cell studies. Nature Communications, 12, 2021.

[20] Shachar Kaufman, Saharon Rosset, Claudia Perlich, and Ori Stitelman. Leakage in data mining. ACM Transactions on Knowledge Discovery from Data, 6:1–21, 2012.

[21] Sayash Kapoor and Arvind Narayanan. Leakage and the reproducibility crisis in machine-learning-based science. Patterns, 4:100804, 2023. PMID 37720327; PMC10499856.

[22] Sean Whalen, Jacob Schreiber, William S. Noble, and Katherine S. Pollard. Navigating the pitfalls of applying machine learning in genomics. Nature Reviews Genetics, 23:169–181, 2021. PMID 34837041.

[23] Eirini Arvaniti and Manfred Claassen. Sensitive detection of rare disease-associated cell subsets via representation learning. Nature Communications, 8:14825, 2017.

[24] Bryan He, Matthew Thomson, Meena Subramaniam, Richard Perez, Chun Jimmie Ye, and James Zou. CloudPred: Predicting Patient Phenotypes From Single-cell RNA-seq. Pacific Symposium on Biocomputing, 27:337–348, 2022.

[25] Travis S. Johnson, Christina Y. Yu, Zhi Huang, Siwen Xu, Tongxin Wang, Chuanpeng Dong, Wei Shao, Mohammad Abu Zaid, Xiaoqing Huang, Yijie Wang, Christopher Bartlett, Yan Zhang, Brian A. Walker, Yunlong Liu, Kun Huang, and Jie Zhang. Diagnostic Evidence GAuge of Single cells (DEGAS): a flexible deep transfer learning framework for prioritizing cells in relation to disease. Genome Medicine, 14:11, 2022.

[26] Guangzhi Xiong, Stefan Bekiranov, and Aidong Zhang. ProtoCell4P: an explainable prototype-based neural network for patient classification using single-cell RNA-seq. Bioinformatics, 39(8):btad493, 2023.

[27] Yuzhen Mao, Yen-Yi Lin, Nelson K. Y. Wong, Stanislav Volik, Funda Sar, Colin Collins, and Martin Ester. Phenotype prediction from single-cell RNA-seq data using attention-based neural networks. Bioinformatics, 40(2):btae067, 2024.

[28] Anastasia Litinetskaya, Soroor Hediyeh-zadeh, Amir Ali Moinfar, Mohammad Lotfollahi, and Fabian J. Theis. Weakly supervised learning uncovers phenotypic signatures in single-cell data, 2024. Preprint at 10.1101/2024.07.29.605625.

[29] Chau Do and Harri Lähdesmäki. Incorporating hierarchical information into multiple instance learning for patient phenotype prediction with single-cell RNA-sequencing data. Bioinformatics, 41(Supplement 1):i96–i104, 2025.

[30] Kyeonghun Jeong, Jinwook Choi, and Kwangsoo Kim. scMILD: Single-cell multiple instance learning for sample classification and associated subpopulation discovery. iScience, 29:115284, 2026.

[31] T. Verlaan, G. A. Bouland, A. Mahfouz, and M. J. T. Reinders. scAGG: Sample-level embedding and classification of Alzheimer’s disease from single-nucleus data. Computational and Structural Biotechnology Journal, 27:3753–3761, 2025.

[32] Felix Fischer, David S. Fischer, Roman Mukhin, Andrey Isaev, Evan Biederstedt, Alexandra-Chloé Villani, and Fabian J. Theis. scTab: Scaling cross-tissue single-cell annotation models. Nature Communications, 15, 2024. PMID 39098889; PMC11298532.

[33] Fabian Pedregosa, Gaël Varoquaux, Alexandre Gramfort, Vincent Michel, Bertrand Thirion, Olivier Grisel, Mathieu Blondel, Peter Prettenhofer, Ron Weiss, Vincent Dubourg, Jake Vanderplas, Alexandre Passos, David Cournapeau, Matthieu Brucher, Matthieu Perrot, and Édouard Duchesnay. Scikit-learn: Machine Learning in Python. Journal of Machine Learning Research, 12:2825–2830, 2011.

[34] Kasia Z. Kedzierska, Lorin Crawford, Ava P. Amini, and Alex X. Lu. Zero-shot evaluation reveals limitations of single-cell foundation models. Genome Biology, 26, 2025. PMID 40251685; PMC12007350.

[35] Constantin Ahlmann-Eltze, Wolfgang Huber, and Simon Anders. Deep-learning-based gene perturbation effect prediction does not yet outperform simple linear baselines. Nature Methods, 22:1657–1661, 2025. PMID 40759747; PMC12328236.

[36] Caleb Weinreb, Alejo Rodriguez-Fraticelli, Fernando D. Camargo, and Allon M. Klein. Lineage tracing on transcriptional landscapes links state to fate during differentiation. Science, 367, 2020. PMID 31974159; PMC7608074.

[37] Kunal Jindal, Mohd Tayyab Adil, Naoto Yamaguchi, Xue Yang, Helen C. Wang, Kenji Kamimoto, Guillermo C. Rivera-Gonzalez, and Samantha A. Morris. Single-cell lineage capture across genomic modalities with CellTag-multi reveals fate-specific gene regulatory changes. Nature Biotechnology, 42:946–959, 2024. PMID 37749269; PMC11180607.

[38] Yaara Oren, Michael Tsabar, Michael S. Cuoco, Liat Amir-Zilberstein, Heidie F. Cabanos, Jan-Christian Hütter, Bomiao Hu, Pratiksha I. Thakore, Marcin Tabaka, Charles P. Fulco, William Colgan, Brandon M. Cuevas, Sara A. Hurvitz, Dennis J. Slamon, et al. Cycling cancer persister cells arise from lineages with distinct programs. Nature, 596:576–582, 2021. PMID 34381210; PMC9209846.

[39] Matthew J. Hangauer, Vasanthi S. Viswanathan, Matthew J. Ryan, Dhruv Bole, John K. Eaton, Alexandre Matov, Jacqueline Galeas, Harshil D. Dhruv, Michael E. Berens, Stuart L. Schreiber, Frank McCormick, and Michael T. McManus. Drug-tolerant persister cancer cells are vulnerable to GPX4 inhibition. Nature, 551:247–250, 2017.

[40] Sydney M. Shaffer, Margaret C. Dunagin, Stefan R. Torborg, Eduardo A. Torre, Benjamin Emert, Clemens Krepler, Marilda Beqiri, Katrin Sproesser, Patricia A. Brafford, Min Xiao, Elliott Eggan, Ioannis N. Anastopoulos, et al. Rare cell variability and drug-induced reprogramming as a mode of cancer drug resistance. Nature, 546:431–435, 2017.

[41] Allen Institute for Brain Science. Seattle Alzheimer’s disease brain cell atlas (SEA-AD). CZ CELLxGENE Discover, dorsolateral prefrontal cortex and middle temporal gyrus microglia, 2024. Dataset identifiers a3198428-dd16-4344-8564-3897c1ccdb4b and c66e3198-c766-499e-a609-7b462a41295b.

[42] Trevor Hastie, Robert Tibshirani, and Jerome Friedman. The Elements of Statistical Learning: Data Mining, Inference, and Prediction. Springer, New York, 2nd edition, 2009.

[43] Adam Paszke, Sam Gross, Francisco Massa, Adam Lerer, James Bradbury, Gregory Chanan, Trevor Killeen, Zeming Lin, Natalia Gimelshein, Luca Antiga, Alban Desmaison, Andreas Köpf, Edward Yang, Zachary DeVito, Martin Raison, Alykhan Tejani, Sasank Chilamkurthy, Benoit Steiner, Lu Fang, Junjie Bai, and Soumith Chintala. PyTorch: An Imperative Style, High-Performance Deep Learning Library. In Advances in Neural Information Processing Systems, volume 32, pages 8024–8035, 2019.

[44] Tamim Abdelaal, Lieke Michielsen, Davy Cats, Dylan Hoogduin, Hailiang Mei, Marcel J. T. Rein-ders, and Ahmed Mahfouz. A comparison of automatic cell identification methods for single-cell RNA sequencing data. Genome Biology, 20:194, 2019.

[45] Brent A. Biddy, Wenjun Kong, Kenji Kamimoto, Chuner Guo, Sarah E. Waye, Tao Sun, and Saman-tha A. Morris. Single-cell mapping of lineage and identity in direct reprogramming. Nature, 564:219–224, 2018. PMID 30518857; PMC6635140.

[46] Malte D Luecken and Fabian J Theis. Current best practices in single-cell RNA-seq analysis: a tutorial. Molecular Systems Biology, 15, 2019.

[47] Volker Bergen, Marius Lange, Stefan Peidli, F. Alexander Wolf, and Fabian J. Theis. Generalizing RNA velocity to transient cell states through dynamical modeling. Nature Biotechnology, 38:1408–1414, 2020.

